# The good, the BATV, and the ugly - first report of BATV vector competence in *Culex quinquefasciatus* from the Southern United States

**DOI:** 10.1101/2024.05.23.595596

**Authors:** Samantha D Clark, Erik A Turner, Jordan M Vivien, Grace B. Buras, Rebecca C. Christofferson

## Abstract

Batai virus is an arbovirus with wide geographic, host, and climactic ranges. BATV infects primarily avian species but can cause disease in ruminants and humans. Louisiana is at particular risk for overlaps in natural and agricultural systems as the Mississippi Flyway passes through the state. We assessed the vector competence of Louisiana *Cx. quinquefasciatus* for BATV and found infection and dissemination rates of 22% and 11.1% at 7 days post-exposure (dpe), and 15.8% and 5.3% at 21dpe. The current H5N1 avian influenza outbreak in dairy cows demonstrates the importance of understanding the overlap in avian and other vertebrate species to inform public health and agricultural biosecurity. Results indicate a moderate vector competence of regional Louisiana *Cx. quinquefasciatus* for BATV. This study presents the first known report of vector competence of BATV in US mosquitoes and establishes the non-zero risk of its emergence in the southern US.

## Background

Batai virus (BATV, *Orthobunyavirus Batai orthobunyavirus)* is an emerging arthropod-borne virus (arbovirus) with an extensive geographic, host, and climactic range of the genus *Orthobunyavirus*, family *Peribunyaviridae*. It is a single-stranded, negative-sense RNA virus with tri-segmented genome consisting of small, medium, and large segments. Segments encode the nucleocapsid, envelope glycoprotein, and RNA-dependent RNA Polymerase respectively (*1, 2*). Since its initial discovery in Malaysia in *Culex gelidus* in 1955, BATV has been detected in several countries across four continents, spanning temperate and tropical climates. It is considered endemic in Europe, Asia, and Africa with a newly emerging strain in Australia (*1*). Previously identified synonymous strains include Calovo virus, Chittoor virus, and recently, a strain of Ilesha virus was re-classified as BATV (*3*). BATV is thought to persist in a bird-mosquito enzootic cycle, and has been isolated from *Anopheles, Culex, Aedes, Coquillettidia*, and *Culiseta* mosquitoes and birds of the orders Passeriformes, Anseriformes, Galliformes, and Gruiformes (*1, 4*). Bovines may also play a key role in transmission as a reservoir or amplification host with moderate seroprevalence across Europe and in China (*5*). Additional epizootic transmission has been observed within ruminants such as deer, goats, sheep, and buffalo, as well as swine, camels, wild hare, horses, donkeys, mules, harbor seals, and humans (*1, 6*).

BATV presents as asymptomatic to mild influenza-like disease in humans, including primarily febrile symptoms of fever, malaise, and myalgia. Reported cases of bronchopneumonia, pleurisy, tonsillitis, gastritis, and malaria-like disease with hospitalizations have occurred (*1, 2, 7*). BATV has been associated with febrile illness and neurological sequelae (*6, 8*). BATV has also been associated with mild egg drop in ducks in China, and was the etiological agent of meningoencephalomyelitis in two captive harbor seals (*9, 10*).

BATV is well characterized with respect to its European distribution, but its potential vector range still remains widely unclassified. Evaluating the breadth of vector species is crucial to determining true distribution, especially in places it has yet to emerge. Without knowledge of competent vector species, the full extent for BATV to become a public health issue is underestimated. In particular, studies evaluating the vector competence of mosquitoes for BATV are crucial to understanding its potential for emergence and reassortment in new geographic regions.

*Culex quinquefasciatus* (‘quinqs’) is a vector of several pathogens affecting humans and animals. It is the primary vector of St. Louis Encephalitis Virus (SLEV), and is a competent vector for West Nile Virus (WNV), Japanese Encephalitis Virus (JEV), Western Equine Encephalitis virus (WEEV), and Sindbis virus (SINV) (*11, 12*). Recently, *Cx. Quinquefasciatus* was implicated in transmission of Chikungunya virus (CHIKV) (*13*). Additionally, ‘quinqs’ have been indicated as an experimental vector of certain strains of Zika virus (ZIKV) (*14*), Usutu virus (USUV) (*15*), Ingwavuma virus (INGV) (*16*), Oropouche virus (OROV) (*17*), Barmah Forest virus, and Ross River virus (*18*). *Cx. quinquefasciatus* was also indicated in transmission of other Orthobunyaviruses in the epizootic outbreak of Rift Valley Fever Virus (RVFV) in Egypt (*11*), and of BUNV in Argentina (*19*). *Cx. quinquefasciatus* is ornithophilic, but will feed on mixed populations of dogs, monkeys, ruminants, equids, and humans opportunistically (*11, 14*). This species is subtropical, with a global distribution in North America, South America, Australia, Asia, and Africa. In the southern United States, this species ranges from the Gulf South – including Louisiana – to southern Iowa (*20*). In this region, *Cx. quinquefasciatus* is the primary vector of WNV and SLEV, which share a common avian-mosquito cycle in the US (*20*). Furthermore, the Flaviviruses USUV and JEV, Alphaviruses WEEV and SINV, and Orthobunyaviruses OROV and INGV have all shown at least laboratory competence within *Cx. quinquefasciatus*, and can all persist and amplify in similar susceptible avian hosts – namely, migratory and wild birds of the orders Passeriformes, Gruiformes, and Anseriformes (*15-17, 21-23*). Indeed, it is this commonality of avian vertebrate hosts that implicates *Cx. quinquefasciatus* as a target for investigating vector competence for BATV.

While BATV has yet to reach the Americas, its transmission ecology is not dissimilar to that of WNV, as is evidenced by an overlap in Europe, and parts of Asia, Australia, and Africa (*1-3, 6, 24-31*), where WNV is maintained in a primarily wild bird-mosquito cycle with vectors in the *Culex* genus (*1, 25*). WNV emerged in North America, and subsequently became established in an avian-mosquito cycle primarily with *Cx. pipiens, Cx. tarsalis*, and *Cx. quinquefasciatus* (*32, 33*), with *Cx. quinquefasciatus* as the primary vector in the southern United States (*20*). BATV also has sufficient overlap with JEV in Asia (*25, 34*). Recent emergence in Australia has heightened concern for JEV emergence in the US and EU, and reports of JEV competence in North American *Cx. quinquefasciatus* already exist (*34-36*). BATV already boasts the largest geographic range of the Orthobunyaviruses, but its potential to emerge in the United States has not been evaluated. Despite its ornithophilic nature, and overlap with WNV ecology, there are no current studies assessing the vector competence of Cx. quinquefasciatus in the US for the transmission of BATV, and only one publication globally assessing vector competence of *Cx. quinquefasciatus* (*6*). In this study, the competence of wild**-**caught ‘quinqs’ from Southern Louisiana was investigated for BATV via infection and dissemination rates, to help assess the public health risk for BATV in the southern United States.

## Methods

### Cell and Virus Strain

African green monkey kidney (Vero) cells (ATCC) were cultured (*37*). BATV (strain MM2222) was originally isolated from Cx. spp. in Malaysia in 1955 (GenBank accession numbers: AB257766, JX8446595-97, and X73464) and was obtained from World Reference Center for Emerging Arboviruses. Passage history at the time of experiment was suckling mouse brain 3 passages, Vero cells 7 passages. Virus utilized for mosquito exposure was freshly collected at 100% CPE at 7dpi the morning of exposure. Infectious titer was 7.3 x 10^5^ pfu/mL determined via Crystal Violet Plaque Assay on Vero cell line prior to experimentation as in (*38*).

### Mosquito Rearing

*Culex quinquefasciatus* larvae were collected using a mosquito emergence chambers in Prairieville, Louisiana and were transported to the ACL2 insectary at LSU and reared in environmental chambers under constant conditions with 18:6h light:dark photoperiods, approximately 80% humidity (monitored via Kestrel LiNK Kestrel DROP 2 devices), and 28°C constant temperature as in (*39*). Mosquitoes were offered 10% sucrose water solution *ad libitum*, but were sugar-starved 48h prior to blood feeding.

### Mosquito Oral Exposure

Adult mosquitoes 10-11 days post-emergence were offered an infectious blood meal comprised of 2:1 bovine blood in Alsevers (Hemostat Labs, Dixon, CA, USA) to infectious supernatant BATV SM3V7. A 3-inch piece of sausage casing was cut and rinsed well with tap water. One end was knotted and filled with water to check for leaks. It was then filled with the pre-warmed blood-supernatant mix and tied off at the other end. In addition, blood-soaked pledgets were placed atop the cage for maximum feeding success. *Cx. quinquefasciatus* were orally exposed to BATV for 2.5 hours. Post bloodmeal, mosquitoes were aspirated and cold anesthetized, sorted, and placed into clean cartons. Mosquitoes were sorted based on visualized blood in the abdomen. Two males were noted to have blood fed and were sorted and kept, as well.

### Sample Collection and Virus Detection

At 7, 14, and 21 days post exposure (dpe) samples of females were taken and wings + legs, and bodies were placed into separate locking Eppendorf tubes with 900uL BA-1 media and Zinc-covered BB’s. Samples were processed as in (*39*). RNA was extracted using the MagMax Viral RNA Isolation Extraction kit per manufacturer instructions (Applied Biosystems, AM1939) then quantified via qRT-PCR on a Roche Lightcycler 96 using Quantabio Ultratough Mastermix (*40*). Primers targeted the BATV M segment described in (*37*). qRT-PCR was performed on each sample in triplicate and detection (yes/no) was determined if at least one triplicate was positive. Quantification of titers was assessed as an average of positives per sample. Prior to experiments, qRT-PCR sensitivity experiments were conducted with a known titer of BATV as described previously in (*40*).

### Statistical Analysis

Observation of infection was calculated as the number of infected female bodies divided by the total number of females tested. Dissemination was likewise determined as the number of positive wing/leg samples divided by the total number tested. Dissemination efficiency was calculated as the total number of positive wing/leg samples divided by the total number of positive bodies. To test for differences in proportion positive at each timepoint, we employed a Chi-square test for multiple proportions (prop.test function in R). Titer per day and across time points was compared via the Mann-Whitney U non-parametric t-test. For the sensitivity assay, difference in detection ability based on sex was determined via a Kruskal-Wallis non-parametric analysis of variance. Limit of detection (LOD) was determined as described in (*40*) utilizing a qPCR LOD calculation R script (*41*).

## Results

### Sensitivity Assay

The qRT-PCR assay used was sensitive for BATV detection in pooled mosquito samples (LOD = 76.1 copies/mL with a 95% confidence). No difference in viral RNA is noted between males versus females (p > 0.05), therefore, the assay yielded statistically similar sensitivity regardless of sex (p = 0.7139). At titers of 10^1^ and above, detection rates were 96.7% (n=30) for combined sexes, 100% (n = 15) for only male pools, and 93.3% (n = 15) for only female pools with one female pool below the limit of detection. At titers of 10^0^, detection rates were 30% for combined sexes (n = 10).

### Cx. quinquefasciatus are moderately competent for BATV

At 7 dpe, 22.2% of exposed females were infected and 11.1% had a disseminated infection. At 14 dpe, none of the 13 exposed females were detected as positive, while at 21 dpe, 15.8% were infected and 5.3% had disseminated infections. In addition, one of the two male mosquitoes that had taken a blood meal was positive for BATV (whole body).

Dissemination efficiency was relatively high, 50% at 7 dpe and 33% at 21 dpe. Cumulatively, 14% of exposed mosquitoes became infected and 6% developed a disseminated infection, with a cumulative dissemination efficiency of 42.9%.

There was no significant difference between the proportion of mosquitoes that were infected at day 7 versus day 21 dpe; between the proportion of mosquitoes with disseminated infections at day 7 versus day 21 dpe or day 14 versus day 21 dpe; or between the dissemination efficiency at days 7 or 21 dpe (p > 0.05).

### BATV Kinetics in Louisiana Cx. quinquefasciatus

At day 7 dpe, the median concentration of BATV RNA genome equivalent in bodies was 1.062 x 10^3^ (IQR: 6.08 x 10^2^, 2.65 x 10^8^)and 4.921 x 10^3^ (IQR: 3.057 x 10^3^, 6.786 x 10^3^) in wing/leg samples. There is no significant difference between the dissemination and infection titers at day 7 dpe (p = 0.5). At 21 dpe, the concentration was 1.00 x 10^2^ (IQR: 6.1 x 10^1^, 2.05 x 10^2^) in the bodies and the only disseminated infection was at 2.451 x 10^9^. There was no significant difference between dissemination and infection titers at day 21dpe (p = 0.8). There is no significant difference in positive infection titers at day 7 dpe vs day 21 dpe (p = 0.057). There is no significant difference in positive dissemination titers at day 7 dpe vs day 21 dpe (p = 0.67).

## Discussion

This is the first demonstration of competence of United States mosquitoes for BATV. While competence was moderate, these data demonstrate the non-negligible potential for BATV to emerge. Additionally, we demonstrate a BATV qRT-PCR assay as a potential tool in mosquito surveillance programs, with a successful detection rate of 50 genomic copies in 96.7% of pools tested. CHITV, the Indian strain of BATV, yields titers within mosquitoes of 5-6 log_10_TCID_50_/mL (*6*) in laboratory settings, suggesting the assay is sensitive for a surveillance setting.

Interestingly, the *Cx. quinquefasciatus* larvae were collected from containers during a record-breaking year where Louisiana experienced severe drought and record high temperatures: 32 days occurred with temperatures exceeding 100 °F, and June-August produced the highest average temperatures ever seen (*42*). The Louisiana Governor declared a State of Emergency as lengthy droughts and wildfires impacted safe drinking water availability across the state (*43*). Generally, *Cx. quinquefasciatus* prefer to lay eggs in stagnant, nutrient-rich water, but will utilize other containers such as tires or pots if other sources of standing water are not available. This is a phenomenon known as water source crowding (*20, 44*). Such an occurrence was also observed during the 2012 drought followed by a record outbreak of WNV in Houston, TX (*44, 45*).

The current H5N1 avian influenza outbreak in dairy cows (*46*) has demonstrated the importance of understanding the overlap in avian and other vertebrate species to inform public health and agricultural biosecurity practices. Louisiana is particularly at risk for overlaps in natural and agricultural systems, as the Mississippi Flyway passes through a large portion of the state. The Mississippi Flyway has been implicated previously in movement of WNV positive passerine species (*47*), showing potential for other zoonotic arboviruses to proliferate in the region.

Similarly, migratory and wild birds have been indicated previously in BATV transmission to new geographic regions via the African-Eurasian flyway (*1*), and although entrance into the US via migratory birds across the Atlantic is less likely, though not implausible, the significance of the Mississippi, Atlantic, Central, and Pacific flyways in emergence of newly emergent zoonotic arboviruses once in the US cannot be ignored.

Further, game animals are proposed to play a role in the maintenance cycle; BATV antibodies have been found in a few species of deer and wild boars (*48*). Louisiana suffers an overpopulation of feral swine and a thriving hunting industry for deer, both of which are in contact with domestic animals. Moreover, Louisiana has a strong agricultural sector with poultry, dairy and beef cattle, goat and sheep, horse, and swine industries – the market value of which totaled approximately 2.5 billion in 2020 (*49*). The top animal agricultural industry is poultry, followed by cattle, with majority of the state’s producers running cow-calf operations, specifically (*49*). BATV has shown proclivity for morbidity and decreased production in livestock (*8, 9*).

While the US has made strides in developing infrastructure for response to arbovirus emergence, the chain of events that led to WNV emergence in the US is repeatable (*32*). Factors of climate change such as milder winters, warmer summers, droughts, and frequent flooding; increased travel and commerce trade; and urbanization and development will continue to increase geographical ranges of competent vectors and create conditions that are right for novel arboviral emergence in previously unaffected regions (*1, 2, 4, 44*). These data demonstrate the potential risk for BATV emergence, with a wide range of potential effects across human and animal health, as well as economic impacts. This points to a need for future study to define the risk of emerging arboviruses such as BATV in the US, especially as it relates to competent vertebrate hosts, particularly avifauna. Ultimately, understanding the landscape of emergence possibilities can inform biosecurity preparedness plans, especially for the agricultural sector, and provide justification for future studies into control and countermeasures.

| Infection and dissemination of BATV in female Louisiana *Culex quinquefasciatus* |  |  |  |  |
| --- | --- | --- | --- | --- |
| Day post exposure | Infection | Dissemination | Dissemination efficiency | N |
| 7 | 22.2% | 11.1% | 50% | 18 |
| 14 | 0% | 0% | NA | 13 |
| 21 | 15.8% | 5.3% | 33% | 19 |

**Figure:**
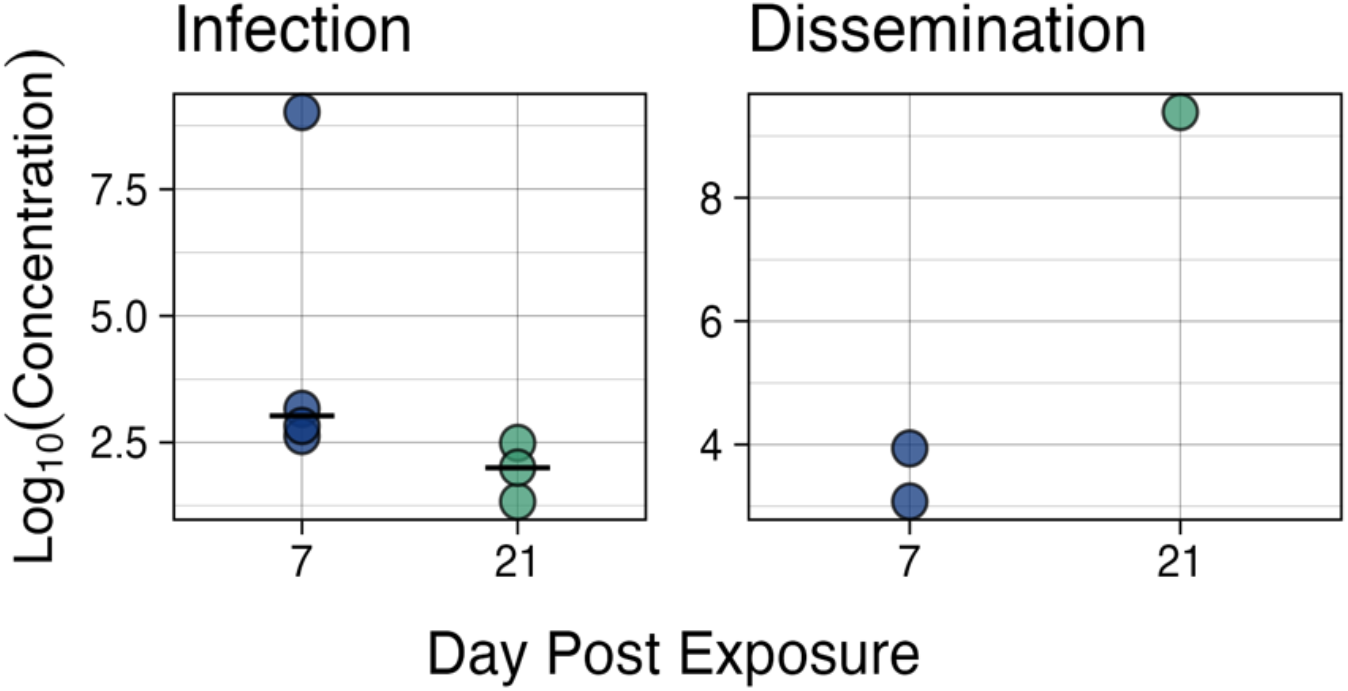
Genome equivalents (log10) of BATV in bodies (infection) and legs/wings (dissemination) of Culex quinquefasciatus at 7 and 21 days post exposure.

